# Transposon Extermination Reveals Their Adaptive Fitness Contribution

**DOI:** 10.1101/2021.11.29.470382

**Authors:** Susanne Cranz-Mileva, Eve Reilly, Noor Chalhoub, Rohan Patel, Tania Atanassova, Weihuan Cao, Christopher Ellison, Mikel Zaratiegui

**Affiliations:** Department of Molecular Biology and Biochemistry, the State University of New Jersey, Piscataway, NJ, USA; Department of Genetics, Rutgers, the State University of New Jersey, Piscataway, NJ, USA

## Abstract

Transposable Elements are molecular parasites that persist in their host genome by generating new copies to outpace natural selection. Here we measure the parameters governing the copy number dynamics of the fission yeast Tf2 retrotransposons, using experimental and natural populations and a strain where all Tf2 copies are removed. Natural population genomes display active and persistent Tf2 colonies, but in the absence of selection mitotic recombination deletes Tf2 elements at rates that far exceed transposition. We show that Tf2 elements provide a fitness contribution to their host by dynamically rewiring the transcriptional response to metabolic stress. Therefore, Tf2 elements exhibit a mutualistic rather than parasitic behavior toward their host.

---

Retrotransposons are Transposable Elements (TE) that can create and integrate new copies through reverse transcription of an RNA intermediate. In this manner, they have colonized most eukaryotic genomes and in some cases constitute the majority of their genetic material^1^. Insertion of new copies in the host genome is expected to result in loss of fitness through mutation of coding and non-coding elements, increased replication and recombination burden, and diversion of cellular resources^2^. Studies in the fly *Drosophila melanogaster* have conclusively shown that the net fitness effect of the TE load in this organism is negative^3–5^. Population genetic surveys also indicate that new insertions are subject to intense negative selection that TE must overcome through active transposition to avert extinction from the host genome^6,7^.

While the selfish character of TE is clear, one possible explanation for their pervasiveness is that they can, at least under some circumstances, provide some fitness benefit to the host genome^8,9^. One proposed mechanism speculates that activation of TE during physiological stress can provide additional genetic plasticity to explore the adaptive landscape, perhaps providing increased evolvability on demand^10,11^. Examples of adaptive transpositions are abundant in the literature^12–14^. However, the question of whether these cases reflect that the persistence of TE is coupled to contributions to host fitness and adaptability or they are simply occasional examples of “hopeful monsters” that would be expected of any mutagenic agent remains unanswered. Regardless of the potential of punctual contributions to host fitness, the clearest evidence that TE are generally selfish elements is that their transposition rates exceed deletion rates, indicating that they are under selective pressure to maintain their presence in the host genome.

TE modulate their effect on host fitness through evolution of insertion target site selection mechanisms. Overall, three different strategies can be identified in the distribution of TE in eukaryotic genomes: (i) widely dispersed random insertions that are then subject to selection; (ii) targeted insertion into dispersed “safe havens” with neutral fitness effect; and (iii) highly directed insertion into specific sites where fitness effects are neutral or even positive^15^. These different strategies are likely coupled to variable transposition rates fine-tuned to counteract the resulting selection pressure.

The fission yeast *Schizosaccharomyces pombe* genomes are exclusively colonized by two highly related Ty3/Gypsy type Long Terminal Repeat (LTR) retrotransposons called Tf1 and Tf2^16^. The genome sequence of the laboratory type strain, derived from the 968/972/975 isolates, has 13 copies of Tf2^17,18^. Two of these Tf2 copies (Tf2-7/8) are present in a tandem repeat that displays copy number polymorphism in laboratory strains derived from 972, forming arrays of up to 5 copies. Tf1 is absent from the type strain but detectable in multiple other isolates^19^. Numerous solo LTR, the product of recombinations that deleted ancestral insertions, indicate a long history of Tf colonization.

Tf1 and Tf2 TE have a clear insertion site preference for type II promoters^18,20^, guided by a tethering interaction between the integrase (INT) and the host-encoded DNA binding factor Sap1^21,22^. This behavior could reflect an adaptation to minimize insertional mutagenesis and its impact on host fitness (a “dispersed safe haven” strategy as detailed above). But a preference for the regulatory regions of protein coding genes suggests that Tf1/2 insertions could modulate host gene expression, with the potential to provide positive fitness by rewiring transcriptional regulatory networks in response to stress challenges that stimulate Tf mobility. Tf1 insertions can act as enhancers to activate transcription of nearby genes, providing a mechanism for this possibility^23^. The potential for adaptive changes was demonstrated in a library of Tf1 insertions generated by overexpression of the transposon followed by selection in CoCl_2_, a heavy metal stressor that upregulates Tf1 transcription^24^. However, not all genes become upregulated by insertion of Tf1 into their promoter,^23^ and the contribution of Tf1 and Tf2 transposons to *S pombe* fitness has not been directly evaluated in the laboratory type strain or in feral isolates.

In order to understand the influence of Tf2 elements on *S pombe* fitness, and how this interaction shapes genome colonization patterns, here we directly measure the parameters driving Tf2 dynamics, transposition and deletion rates as well as selection coefficients, using experimental and natural populations and a laboratory strain in which all Tf2 copies have been deleted.

## Results

### Tf1 and Tf2 copy number dynamics in fission yeast natural isolates

TE element copy number is determined by transposition and recombination processes (Supplementary Figure 1) and natural selection. Like other LTR retrotransposons, Tf2 elements can transpose by INT-mediated insertion or by Gene Conversion (GC)^25^. Recombination between the LTR can lead to deletion through Intra Chromatid Recombination (ICR) or Unequal Strand Chromatic Exchange (USCE). We sought to measure the relative contribution of transposition and deletion processes in the TE colonies of natural fission yeast isolates. A recent survey of natural fission yeast isolates from multiple sources revealed wide polymorphism in the number and location of Tf insertions, as assessed by the presence of LTR in short-read sequencing data^26^. However, this study did not evaluate the presence of CDS that would indicate the copy number of full-length insertions nor made a distinction between Tf1 and Tf2. We analyzed the short-read sequence generated by this study to assess the presence and activity of Tf2 and Tf1 colonies in natural fission yeast isolates.

We first measured the copy number of Tf1 and Tf2 CDS using the sequence coverage over the *gag* gene, which is highly divergent between Tf1 and Tf2^27^, normalized to the coverage of multiple single copy essential genes of the same size and AT content ^28^ (Supplementary Figure 2). This analysis revealed that Tf2 and Tf1 CDS show a wide variability in copy number ranging from 0 to more than 100 copies (Figure 1a). While Tf2 is almost ubiquitous, Tf1 appears to have been lost from some clades, including the one that contains the 972 laboratory strain.

**Figure 1:**
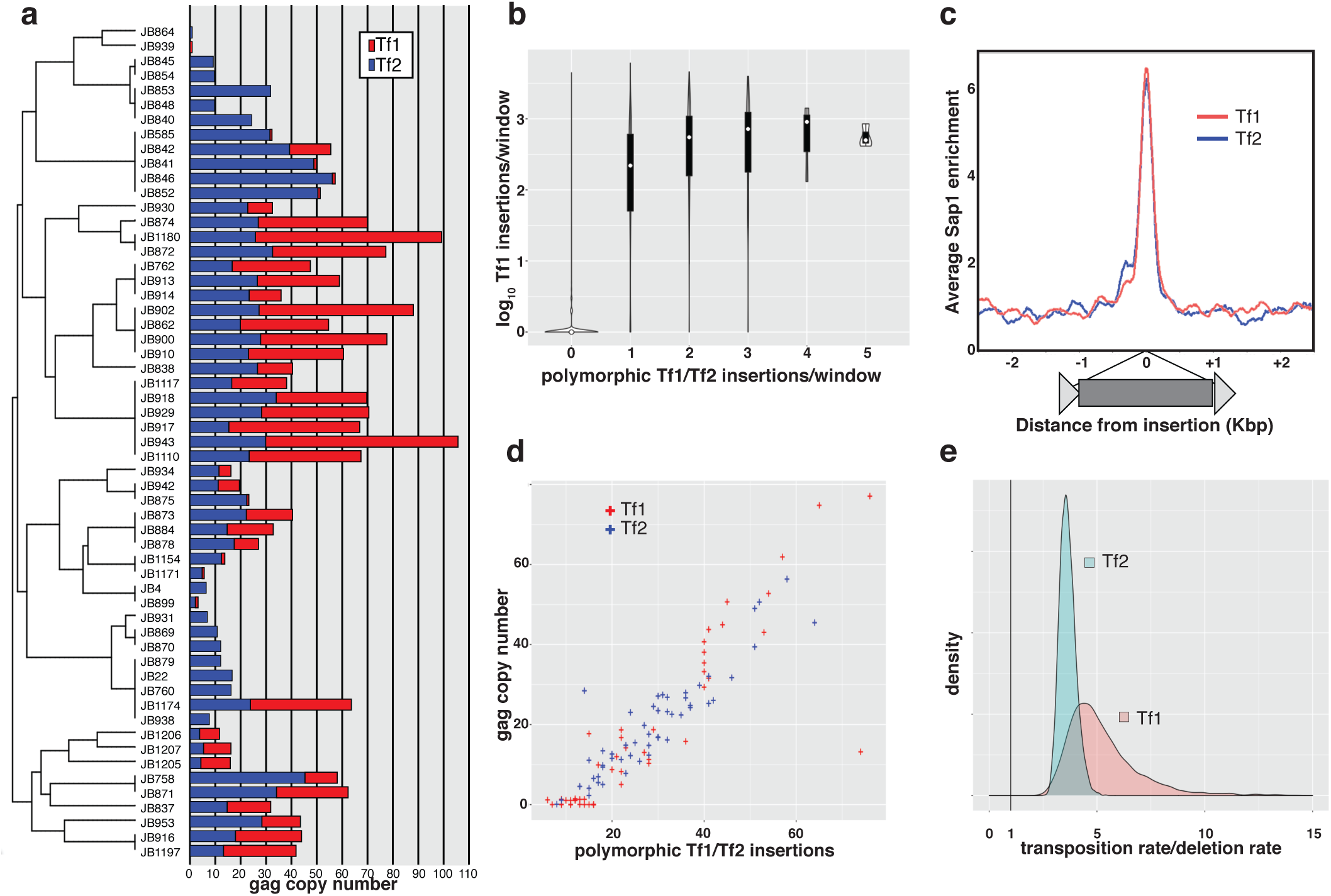
Tf1 and Tf2 copy number dynamics in natural fission yeast isolates. a. Estimated copy number of Tf1 (red bars) and Tf2 (blue bars) in the natural isolates arranged in the phylogeny tree described in ^26^. b. Genomewide correlation between the insertion profile of overexpressed Tf1 ^21^ and the number of polymorphic Tf1 and Tf2 insertions detected in the natural isolates, in 50bp windows. c. Sap1 enrichment around polymorphic Tf1 (red) and Tf2 (blue) insertions. d. Scatterplot for Tf1 (red) and Tf2 (blue) *gag* copy number and polymorphic insertion copy number. e. Density plot of posterior estimates for transposition/deletion rate ratios in Tf1 (red) and Tf2 (blue).

We next analyzed the number and position of Tf1 and Tf2 insertions using the MELT pipeline^29^ followed by assembly of mini-contigs and BLAST alignment. This analysis revealed a significant undercount of the previously reported polymorphic insertions^26^ due to frequent clustering of insertion sites within small target windows. As a result, we counted 1376 polymorphic insertions of Tf1 and Tf2 combined, a 60% increase over the original count (Supplementary Table 1 and Supplementary Figure 2). Mini-contig assembly permitted the identification of Target Sie Duplications (TSD) in most of these insertions. In addition, analysis of genome-wide insertion counts in 50bp windows (Figure 1b) revealed that the polymorphic insertions were concentrated in sites that exhibit Tf1 insertion hotspot activity^21,22^ and were placed within a peak of Sap1 enrichment^30^ (Figure 1c). We conclude that the polymorphic insertions detected in our analysis correspond to canonical Integrase-mediated and Sap1 guided insertion. Tf1 and Tf2 *gag* copy number correlated very strongly with the number of polymorphic insertions within each strain, providing a linear relationship between the number of full insertions and the number of total insertions including solo LTR resulting from deletion events (Figure 1d). Fitting a simple transposition-deletion model that assumes neutral fitness and constant transposition and deletion rates, the observed distribution of polymorphic LTR and *gag* copy number indicates that transposition rates apparently exceed deletion rates by between 3 fold (Tf2) and 5 fold (Tf1) (Figure 1e).

### Tf2 deletion rates exceed transposition rates

Since the observed distribution of Tf2 colonies in natural isolates does not directly provide a measurement of transposition and deletion rates we sought to measure them in controlled conditions. We measured transposition rates using a consensus Tf2 copy marked with a *neoR* antibiotic resistance gene in antisense orientation interrupted by an artificial intron in the sense orientation. In this manner, a mobilization mediated by cDNA confers resistance to the antibiotic G418. To account for GC events we characterized the G418 resistant colonies by PCR to identify mobilizations by GC into pre-existing insertions, whose positions are well known (Figure 2a). We detected a mobilization rate of 5e-8, and a GC frequency of 80%, resulting in a transposition rate of ∼1e-8 mobilizations per generation (Figure 2b). This is in agreement with previously reported transposition rates^31,32^ and with the preference of Tf2 to mobilize by GC^25^.

**Figure 2:**
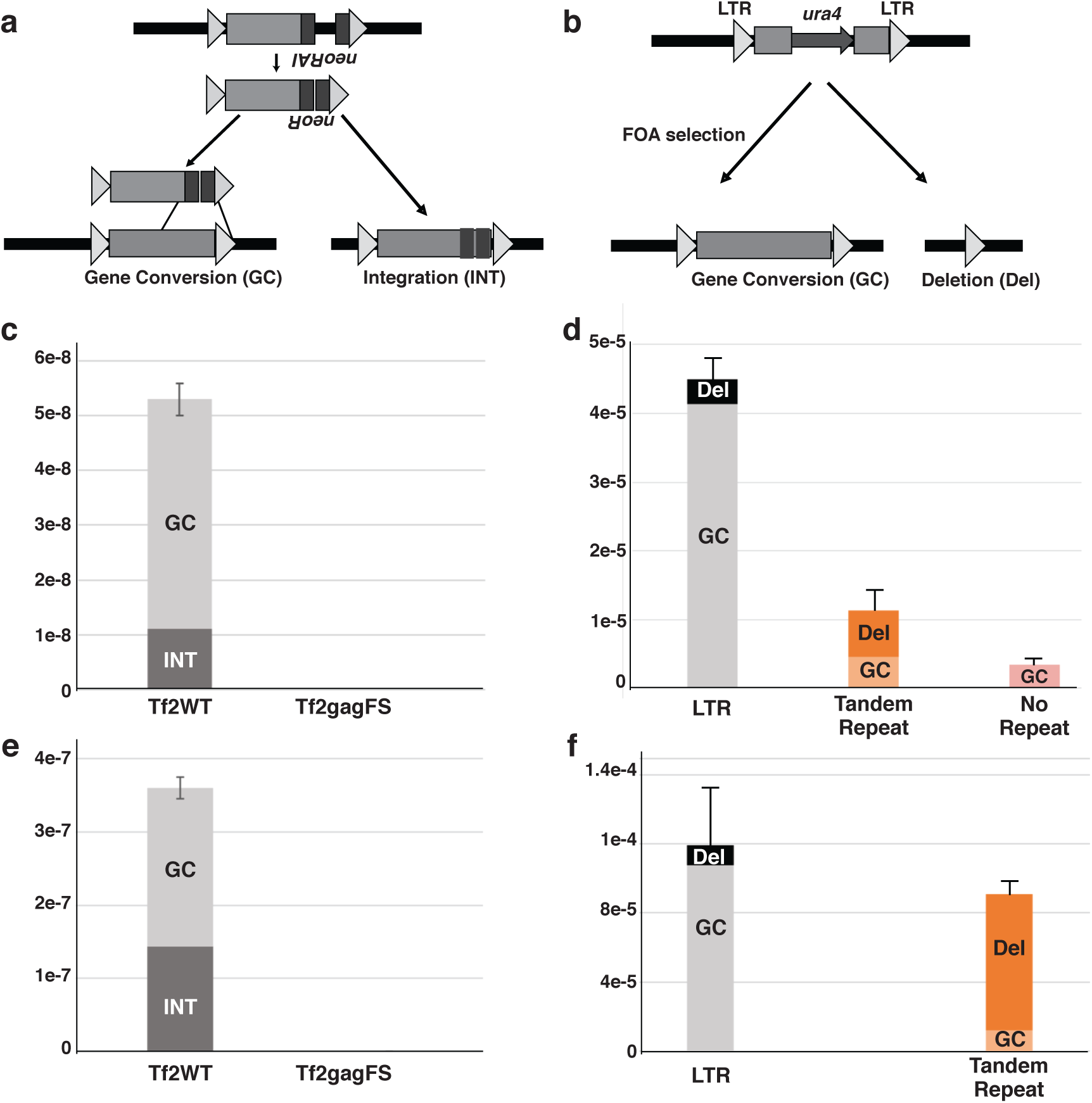
Transposition and recombination rates in genetically marked Tf2 reporters. a: Tf2 transposition reporter. b: Tf2 recombination reporter. c, e: Transposition rates from a WT and *gag* frameshifted Tf2 in mitotic (c) and meiotic (e) cycles. d,f: Recombination rates in a Tf2 element flanked by LTR, a duplication of upstream sequence without the LTR (Tandem Repeat), or unique sequence (No Repeat) in mitotic (d) or meiotic (f) cycles. GC: Gene Conversion. INT: Non-GC mobilization. Del: Deletion. Error bars depict standard error.

In a similar experiment, we measured deletion rates by assaying the loss of a *ura4* genetic marker inserted into the entopic insertion Tf2-6. Loss of *ura4* can be selected by treatment with 5-Fluoroorotic acid (5-FOA). Again, we distinguished deletion events (that lose all the coding sequence) from GC events (that restore the coding sequence from an ectopic template erasing the *ura4* gene) by PCR of the selected colonies (Figure 2c). The overall *ura4* loss rate was 4.5e-5 per generation, with a GC frequency of around 90%, yielding a deletion rate of ∼4e-6 per generation (Figure 2d).

We simultaneously determined if this high deletion rate was due to the presence of the tandemly arranged LTR flanking the coding sequence. Substituting the LTR of Tf2-6 by a tandem repeat generated by duplicating 350bp of upstream sequence (the size of the Tf2 LTR) downstream of the transposon resulted in a similar deletion rate but a lower GC frequency (∼40%) (Figure 2d). This lower GC frequency could be explained by the decreased extension of homology with other Tf2 elements after removal of the LTR. Deletion of both LTR leaving no tandemly repeated sequence resulted in complete loss of deletion events, and all 5-FOA resistant colonies showed to be the result of GC (Figure 2d). Together, these results indicate that deletion events occur at high frequency, exceeding transposition rates, and are an inescapable consequence of the tandemly arranged LTR flanking the coding sequence.

These assays were carried out with asexual mitotically growing cells in essential media rich in glucose and nitrogen, conditions that might not reflect the fission yeast native environment. We hypothesized that growth conditions in the wild could result in higher Tf2 expression and increase transposition rates to exceed deletion rates. In particular, culture in Nitrogen-poor media increases Tf2 expression^33^, and also induces sexual differentiation^34^, leading to mating and meiosis. Considering that deleterious TE require sexual reproduction to maintain their presence in the genome^35^ Tf2 could have evolved a self-regulatory mechanism to restrict its mobilization to meiotic cycles. To evaluate this possibility, we measured the transposition and deletion rates with the same reporters introduced into a homothallic strain grown in malt extract to induce mating and meiosis, followed by spore selection and germination under selective conditions. This experiment revealed that both the transposition and deletion rates increased around ten fold in meiotic cycles, with similar GC frequencies, leaving the relative transposition/deletion rate balance unchanged (Figure 2e,f). We conclude that, in our ectopic models of mobility and recombination, Tf2 deletion rates exceed transposition rates by a large margin.

### Entopic Tf2 dynamics in the absence of selection

While consistent with measurements previously reported, the Tf2 transposition/deletion rates measured in our experimental model may not faithfully represent the parameters of Tf2 activity in a natural setting, as we used Tf2 marked with reporter genes that could affect its activity or stability. Furthermore, since our recombination assay selected for loss of the *ura4* marker it would not reveal recombination events that result in Tf2 tandem duplication (Supplementary Figure 1). We therefore sought to measure the parameters of entopic Tf2 element activity. Entopic TE mobility parameters can be estimated in the absence of natural selection through the characterization of Mutation Accumulation (MA) lines. An MA experiment in *D melanogaster* revealed that TE transposition rates far exceed deletion rates^36^. We undertook a similar analysis on data obtained in two independent MA experiments performed in *S pombe* by random colony picking in a total of 175 lines propagated between 1700 and 1900 generations and then sequenced through short read paired end sequencing^37,38^.

First, we analyzed the sequencing data to look for new insertions with the MELT pipeline^29^. We detected one single new Tf2 insertion at position III:756632 (Figure 3a). This position coincides with a Tf1 insertion hotspot within a Sap1 binding peak^21^. Assembly of a mini-contig around the insertion using reads mapping to that position from the MA line where the insertion was detected revealed a consensus Tf2 LTR and a 5bp TSD. Together, these observations indicate that this insertion was the product of canonical INT mediated mobilization guided by Sap1, yielding a calculated transposition rate of 1.9e-7 per Tf2 per generation.

**Figure 3:**
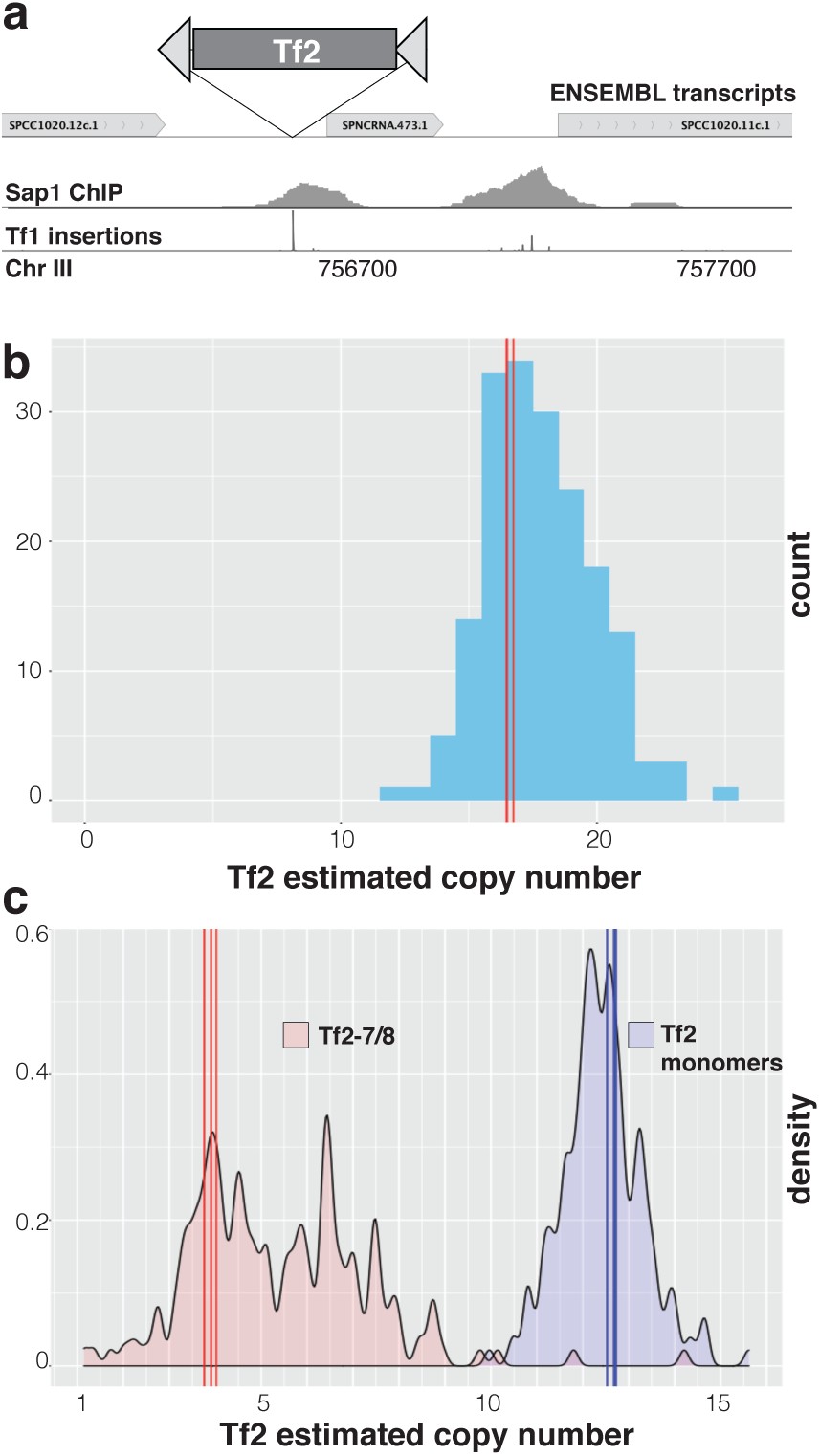
Tf2 copy number dynamics in MA lines. a. Genome environment of the canonical INT-mediated insertion detected in one of the MA lines. Sap1 ChIP from ^30^. Tf1 insertion profile from ^21^. b. Histogram of total Tf2 copy number in the initial strains (red vertical lines) and final MA lines (blue bars). c. Tf2 copy number separated between monomeric Tf2 (blue) and Tf2-7/8 array (red) in the initial strains (vertical lines) and the final MA lines (density plots).

We then analyzed the sequence data for signs of homologous recombination by analyzing copy number and specific polymorphisms in the CDS. First, we quantified the copy number of full-length Tf2 by coverage normalization^28^. This analysis revealed an unexpectedly broad variability in Tf2 copy number in the MA lines. From a starting copy number of 16 in the initial strains, the MA lines ranged from 12 to up to 25 copies per genome, with a trend toward a gain in copy number (mean=+1.2 copies; median=+1.14 copies; sd=2.05; IQR=2.71) (Figure 3b). Since only one new transposition was observed in one of the MA lines, the excess variability must be due to mitotic recombination processes (Supplementary Figure 1).

The frequency of a polymorphism present in all copies of the Tf2-7/8 insertion in the read data allows us to monitor the repeat number of this tandem array. From this information we estimate that the starting strains of the MA experiments had a 4 copy array, consistent with the structure observed in long-read sequencing performed in the 972 strain^39^. Interestingly, the frequency of reads with the Tf2-7/8 polymorphism in the MA lines correlated strongly with the overall copy number (R^2^=0.67, p<1e-6) (Supplementary Figure 3), indicating that the repeat number within the Tf2-7/8 array is the main contributor to total Tf2 copy number variation in the MA lines. If we separate the overall copy number between that attributable to the Tf2-7/8 array and that attributable to the monomeric insertions (Figure 3c), we observe that the size of the Tf2-7/8 array is very polymorphic, with a net tendency to grow in copy number (mean=+1.47 copies; median=+1.30 copies; sd=1.86; IQR=2.44; Wilcoxon test p< 1e-16). In contrast, monomeric Tf2 exhibit a narrower distribution with a small overall loss in copy number (mean=-0.26 copies; median=-0.30 copies; sd=0.88;IQR=1.03; Wilcoxon test p=2.3e-5). These results indicate that the mechanisms governing recombination in multimeric Tf2 insertions are different from those in monomeric insertions: the monomeric insertions conform to the behavior expected from the concerted action of ICR and USCE^40^, but the Tf2-7/8 array exhibits a growth bias that does not fit this model. We estimated the rates of ICR and USCE using the mean and variance of the distribution of monomeric Tf2 copy number in the final MA lines, yielding rates (expressed as events per Tf2 per generation) for ICR of 1.08e-5 (bootstrap SE=2.91e-6) and for USCE of 2.27e-5 (bootstrap SE=4.17e-6), for a total deletion rate of 2.21e-5 (bootstrap SE=2.28e-6). In summary, the entopic Tf2 copy number dynamics on MA lines indicates that in the absence of selection deletion by recombination processes dominate over canonical transposition by two orders of magnitude. This is in agreement with the rates observed with genetically marked transposition and deletion reporters. In addition, we observe that copy number dynamics differ greatly between the Tf2-7/8 array and other Tf2 present in monomeric form.

### Tf2 dynamics in a Tf2-free strain

The discrepancy between transposition/deletion ratios in the natural isolates and the experimental populations of laboratory strains indicates that the assumptions of constant rates and/or neutrality do not hold. Natural selection could distort the apparent transposition/deletion ratio, or transposition rates could decrease with increasing copy number by TE self-regulation^41^. We decided to measure the effect of the transposon colony on transposition/deletion rates and host fitness by removing all entopic Tf2 elements from the laboratory type strain. These Tf-null strains provide a completely isogenic platform to directly test hypotheses about TE/host interactions (Tf-null strain generation is described in the Materials and Methods, Supplementary Figure 4 and Supplementary Table 2).

We measured the transposition rates of genetically marked Tf2 in the presence and absence of the entopic Tf2 colony. These experiments showed that mobilization rates are substantially reduced in the Tf-null strain (from 6e-8 to 1e-8) (Figure 4a). Analysis of the transposed strains shows that this is due to the loss of GC into entopic Tf2 insertions (80% in WT vs 0% in Tf-null strains), while new transpositions remain unchanged (∼1e-8). In agreement with this result, removal of all entopic Tf2 abrogated transposition of an overexpressed Tf2 with an INT frameshift mutation that allows only GC mediated mobility (Figure 4b). The introduction of a *sap1* mutation previously shown to decrease INT-dependent Tf1 mobilization^21^ resulted in complete loss of transpositions in the Tf-null strain (Figure 4b). Together, these results indicate that Tf2 can only mobilize in the Tf-null strain by canonical INT mediated and Sap1-guided insertion, having lost all GC targets^25^. However, the rates of canonical transposition remain unchanged as compared with the WT strain with the full entopic Tf2 colony.

**Figure 4:**
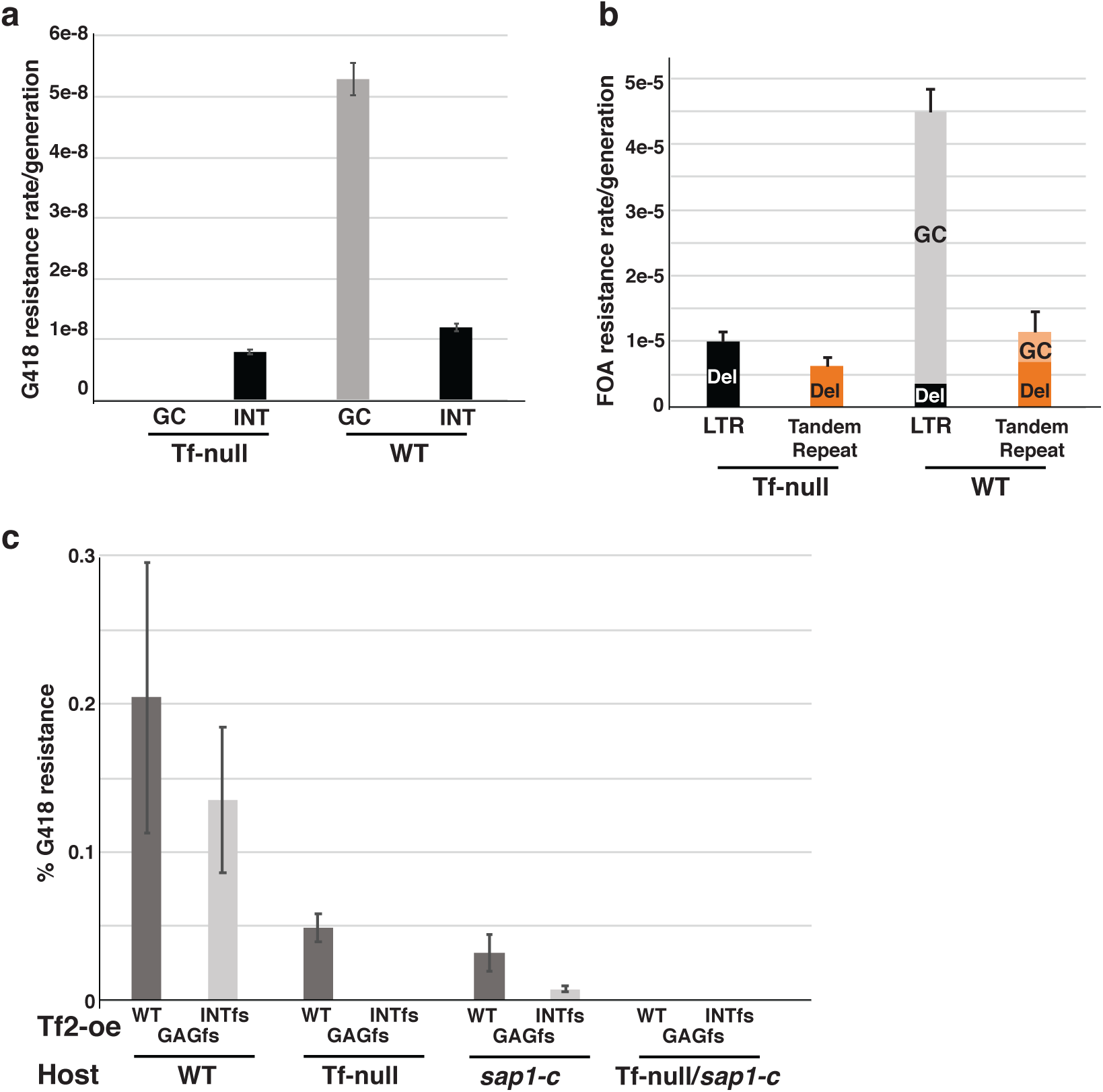
Transposition and deletion of Tf2 in the Tf-null strain. a. Mobilization and b. recombination rates in WT and Tf-null strains using genetically marked reporters as described in figure 1. c. Mobilization frequencies of overexpressed Tf2 (Tf2-oe) vectors with WT sequence (WT), *gag* frameshift (GAGfs) or INT frameshift mutations (INTfs) in strains with (WT) or without (Tf-null) the Tf2 colony, and wild type (WT) or mutated *sap1* (*sap1-c*). Error bars depict standard error.

We next measured the recombination rate in Tf-null strains. Similarly to the transposition experiment, recombination rates drop substantially in the Tf-null strain (4.5e-5 to 1e-5), but only due to the loss of GC events (92% vs 0%) (Figure 4c). These results, while confirming the importance of recombination processes on entopic Tf2 copies, indicate that the rates of deletion and canonical transposition do not depend on Tf2 abundance at least up to the copy number found in the type strain.

### Entopic Tf2 insertions have a positive fitness effect

The availability of isogenic strains differing only in the presence or absence of TE allows for the direct measurement of the effect of the TE colony on host fitness. We carried out a competition experiment to measure this effect. In addition, we evaluated the effect of treatment with CoCl_2_, which mimics hypoxic conditions and activates Tf2 transcription^24^, as well as the effect of mating type to account for the potential effect of expression in h^-^ cells of *mat1-m* encoded factor Mc, a homolog of the sex determination factor SRY that binds to LTR^42^. For this experiment, we mixed equal numbers of cells of the same mating type to prevent sexual reproduction, and passaged them twice a day for ∼140 generations, ensuring the cultures never reached saturation (Figure 5a). We then fit the genotype proportions as a function of relative fitness between the two genotypes, with additive effects of mating type and CoCl_2_ treatment, and number of generations^43^. If the presence of the Tf2 colony confers a negative fitness effect to the host we would observe a relative fitness w_tf0/wt_ >1; if the Tf2 colony brings adaptive value to the host we would observe w_tf0/wt_ <1. Surprisingly the baseline w_tf0/wt_ is 0.9981;[0.9980,0.9983] (median, [HPDI 89%]) (Figure 5b), indicating that the Tf2 colony confers a fitness advantage. The total selection coefficient *s*_Tf2_ is -2e-3, and assuming a purely additive effect of all Tf2 insertions the average !̅_Tf2_ is -1.5e-4 per insertion. Both the h^-^ mating type and CoCl_2_ treatment influenced w_tf0/wt_ with a positive sign (Figure 5c) indicating that Mat2Mc expression and CoCl_2_ treatment decreased the beneficial effect of Tf2 presence on fitness.

**Figure 5:**
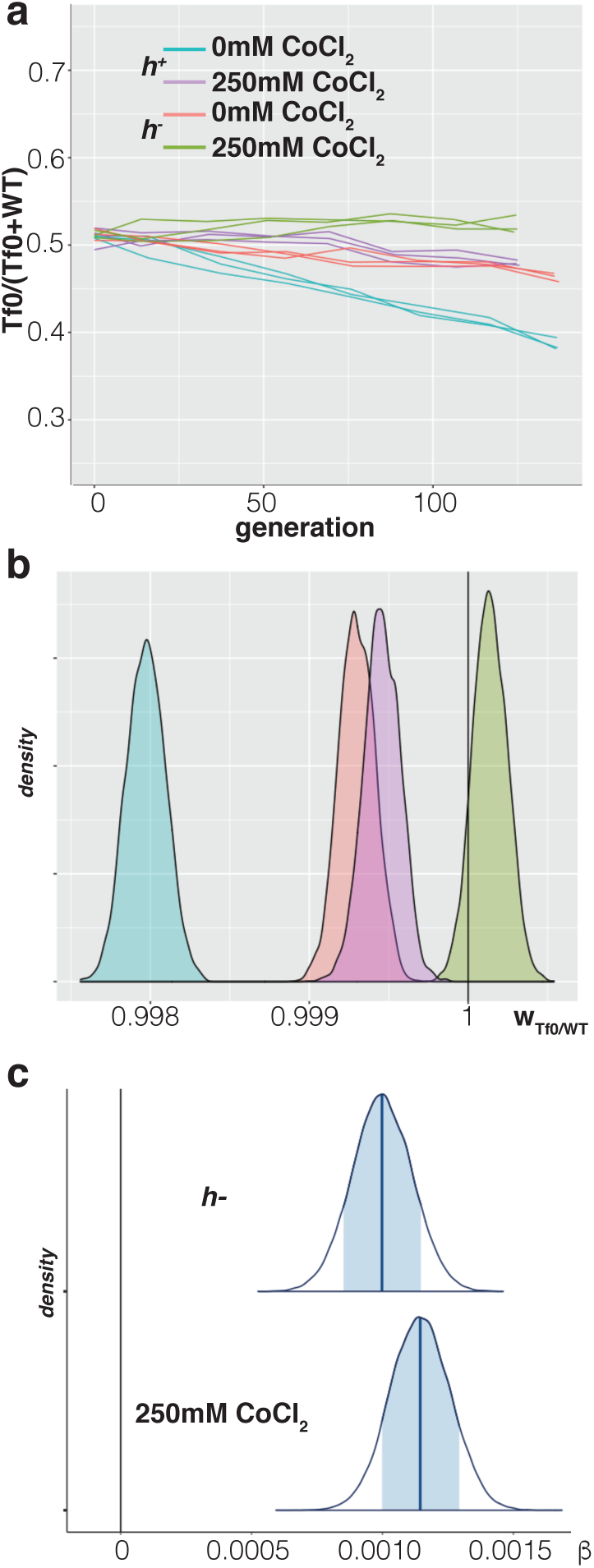
WT versus Tf-null growth competition assays. a. Frequency of the Tf-null specific polymorphism in competition cultures of both mating types (h^+^/h^-^) with and without treatment with CoCl_2_. b. Density plots of posterior probability distributions of relative fitness of Tf-null over WT (w_Tf0/WT_). The vertical line at 1 depicts the expectation of neutrality. c. Density plots of posterior probability distribution of the coefficients for h^-^ and CoCl_2_ treatment. The vertical line at 0 depicts the expectation of no effect of the factor on w_Tf0/WT_.

### Tf2 insertions target ncRNA stress regulons

The related TE Tf1 has shown the potential to provide adaptive value by influencing the expression of stress response genes^23,24^. To investigate the mechanism by which the Tf2 colony increases the fitness of the host genome and CoCl_2_ treatment decrease fitness in the presence of Tf2 we performed RNAseq in WT and Tf-null cells grown with and without CoCl_2_, with a factorial design to measure main effects and interaction. While CoCl_2_ treatment results in widespread expression differences (Figure 6a,c), the absence of Tf2 elements had virtually no effect on gene expression in either condition (no interaction between CoCl_2_/Tf-null detected, Figure 6b,d). Only three genes showed differential expression between the WT and Tf0 strains (Figure 6d). Two of these, *mic10* and the non-coding RNA (ncRNA) *SPNCRNA.1059*, immediately flank the Tf2-7/8 array insertion (Figure 6c). Mic10 is a subunit of the MICOS complex, an organizer of the inner mitochondrial membrane, where it enables respiratory metabolism^44,45^. In fission yeast many respiratory metabolism genes are regulated in *cis* by expression of nearby ncRNA^46,47^. The transcriptional regulation that enables the shift from fermentative to respiratory metabolism upon diauxic shift is mediated by the coordinated action of DNA binding factors Scr1, Tup11 and Rst2^48^. We plotted the binding of Scr1 and Tup11 in cells grown in glucose rich media as well as Rst2 in cells starved for glucose and observed clear peaks for these factors downstream of *SPNCRNA.1059* (Figure 6e). Consistently, both *SPNCRNA.1059* and *SPNCRNA.1058*, an overlapping ncRNA antisense to *mic10*, become upregulated in glucose and sucrose starvation conditions^48^. This arrangement is typical for a regulatory cassette that responds to the diauxic shift, suggesting that the insertion of Tf2-7/8 and subsequent amplification of the array to two or more tandem copies separated *mic10* from its *cis* transcriptional regulators. If the Tf2 insertions in the WT strain confer a competitive advantage over the Tf-null strain by modifying the regulation of metabolic genes, we would expect that the relative fitness would be sensitive to growth conditions. To evaluate this hypothesis we performed a competition assay as before, but allowing the cultures to reach stationary phase passaging them every two days. In this case w_tf0/wt_ is 0.9999;[0.9994,1.0004] (median, [HPDI 89%]), a result consistent with complete neutrality of the Tf2 colony (Figure 6f,g). This indicates that the effect of the Tf2 colony on host fitness depends on growth conditions, further implicating the regulation of metabolic genes.

**Figure 6:**
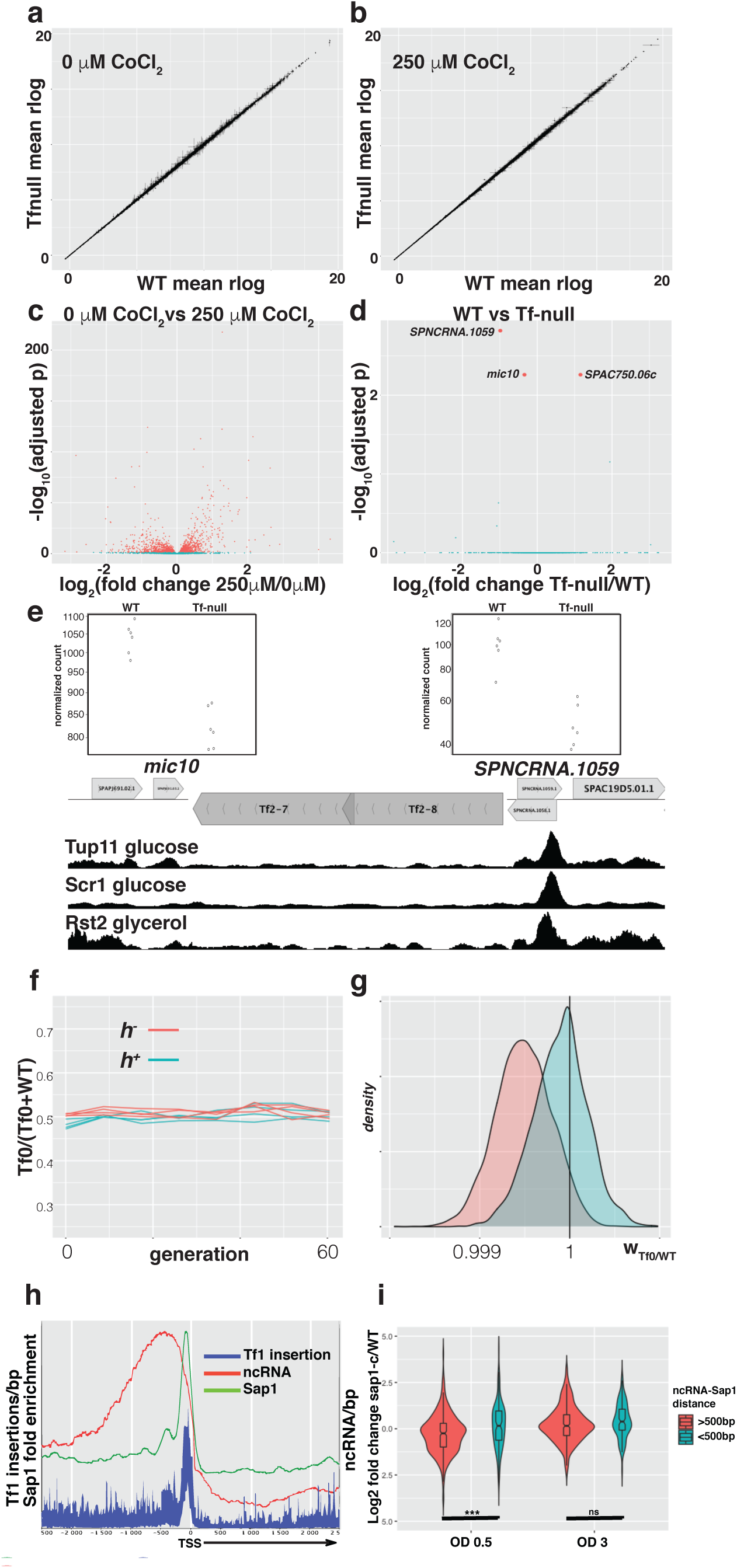
Influence of the Tf2 insertions on host gene expression. a,b. Scatterplot of mean regularized log (rlog) transformed RNAseq read counts of WT and Tf-null strains in (a) 0 μM (b) 250 μM CoCl_2_. c,d. Volcano plots of (c) 0 μM vs 250μM CoCl_2_ and (d) WT vs Tf-null strains RNAseq. The three genes detected as significantly changed in the Tf-null strain are labeled by name. e. Genomic map of the two genes changed in Tf-null, positioned adjacent to the Tf2-7/8 array, with normalized read counts (above) and Tup11, Scr1 and Rst2 enrichment (from ^48^) below. f,g. Growth competition assay between WT and Tf-null strains taken to saturation conditions, as in Figure 6. h. Distribution of Sap1 enrichment (green), Tf1 insertions (blue) and ncRNA (red) around protein coding gene TSS. i. Log2 fold change of intergenic ncRNA expression by RNAseq, classified by their proximity to a Sap1 peak, as measured in early exponential growth (OD=0.5) and early saturation growth (OD=3). (***: p<0.001; ns: p>0.05).

The influence of Tf2 insertions on host fitness through regulation of metabolic genes raises the possibility that the TE has evolved targeting strategies geared toward rewiring ncRNA dependent *cis*-regulatory cassettes. Tf element insertion is guided by Sap1^21,22^ (Figure 6h). *sap1* mutants exhibit a precipitous loss of viability upon reaching stationary phase in rich media^49^, suggesting that its function is important for the response to metabolic stress. We evaluated whether Sap1 is involved in the regulation of intergenic ncRNA by RNAseq of the *sap1-c* mutant growing in early exponential (OD=0.5) and early saturation (OD=3) phases in rich media. This analysis revealed that the *sap1-c* mutant in exponential growth shows upregulation of ncRNA located near Sap1 binding peaks (Figure 6g) and genes involved in the Core Environmental Stress Response (Supplementary Table 3). These results indicate that Sap1 directly represses ncRNA expression during exponential growth. Taken together, these results suggest that Tf1 and Tf2 target ncRNA regulatory networks that respond to changing metabolic needs.

## Discussion

One of the main tenets of TE biology is that if a parasitic (fitness-negative) TE is to maintain a stable presence in the host genome its transposition rates must exceed deletion rates. In this study, we show that the Tf2 element exhibits the opposite strategy, providing adaptive fitness to overcome its high deletion rates. Analyses of ectopic reporters and MA lines confirm that the tandem LTR of monomeric Tf2 elements results in high frequency of recombination events with a net tendency to copy number loss due to the excess of deletion events by ICR over new insertions. Interestingly, both transposition and deletion frequencies increase in conditions that upregulate Tf2 transcription, resulting in an unchanged preponderance of deletions over transpositions. This could be explained if the upregulation in transcription that allows for increased transposition also causes an increase in the genetic insults that are resolved by homologous recombination^50^. A similar phenomenon has been observed in the *S cerevisiae* Ty3 element^51^.

Considering the high recombination rates of Tf2 elements, we speculate that the regulation of recombination of these TE is an important aspect of copy number control that could affect their persistence in the host genome. In this respect, the unexpected growth bias of the Tf2-7/8 array could represent a mechanism that maintains a reservoir of Tf elements resistant to deletion by ICR.

Extrapolating the observed recombination and deletion rates and assuming complete neutrality of Tf2 insertions one would predict that the Tf2 colonies would be completely extinct within ∼3.6e6 generations^40^. On the other hand, population genetics data from natural fission yeast isolates^26^ show that Tf2 TE are actively transposing, creating new insertions that result in highly polymorphic transposon colonies. The contrast between these two observations could be reconciled if we admit that selection pressures acting on natural fission yeast populations drive the increased survival of Tf2 elements.

Using a laboratory type strain with all Tf2 removed, we can observe that the Tf2 colony does provide a net fitness advantage to its host. While the fitness effect is relatively strong once the effective population size of fission yeast^38,52^ is considered (N_e_s∼2e4), it is also highly dependent on growth conditions, being effectively neutral upon growth to saturation. This could indicate that the fitness effects of insertions present in a natural isolate are transient and highly dependent on changing environmental conditions. As a result, surveys of Tf colonies in natural isolates might show no signs of positive selection (Supplementary Figure 2), reflecting instead recent transposition activity and population structure^53^. The insertional preference of Tf1 and Tf2 for the promoters of protein coding genes provides a potential mechanism for the fitness contribution of TE insertion. Tf1 can provide an enhancer activity increasing the expression of nearby genes^23,24^. However this activity is not universal, as not all promoter insertions result in transcriptional changes. Consistently, in our analysis of all the Tf2 insertions present in the type strain we could only observe changes in the expression of genes located next to the Tf2-7/8 array. The original insertion that later expanded to an array would have separated the *mic10* gene, involved in respiratory metabolism, from two ncRNA likely driven by the Scr1/Tup11/Rst2 transcription factors that orchestrate transcriptional responses to changing carbon source conditions^48^. It is possible that the Tf2-7/8 array is the only insertion providing a fitness benefit to the laboratory type strain, and that its tandem structure is the result of selection acting on the outcome of USCE on this insertion.

The fact that the positive fitness effect of the Tf2 colony fades when growth is driven to saturation supports the potential involvement of the metabolic needs of the host. Fission yeast natural isolates are closely associated to human activities involving fermentation^54^, where an initially glucose-rich environment supports rapid growth until glucose is exhausted and other carbon sources must be used^55^. In fission yeast the transcriptional regulatory changes that accompany this change, termed the diauxic shift, are often carried out by ncRNA that control protein coding genes in cis^46,56^. We have shown that Sap1, the DNA binding factor that guides Tf1 and Tf2 elements to insert on type II promoters, regulates ncRNA expression near genes associated with the core stress response pathways. Tf1 and Tf2 could alter the regulation of genes involved in the diauxic shift by severing their association with cis-regulatory ncRNA^57^. Indeed, the Sap1 binding region placed in the NFR of protein coding genes is ideally placed to guide such mutations, as ncRNA are commonly located upstream of core promoters (Figure 6a). It is worth noting that genes whose expression is affected by Tf1 insertion were very often ncRNA^23^. While these insertional mutations may provide a positive fitness effect in specific growth conditions, they nevertheless constitute a loss of complexity that can come with evolutionary tradeoffs. The regulation of the diauxic shift is itself an evolvable trait that can drive the evolution of carbon source generalist and specialist variants, but any such commitments are made at the expense of fitness in some conditions^58^.

Since ncRNA show no coding potential the change induced by Tf1 or Tf2 insertion may be completely or partially reversible through deletion by ICR^57^. In this manner, the regulation of metabolic and stress response genes may be rapidly and dynamically altered by TE element activity responding to changing environmental conditions. Traces of repeated insertion and deletion are visible in the fission yeast genome, where some gene promoters exhibit multiple LTR remnants of ancestral insertions, and independent insertions into the same promoters, sometimes in the exact same position, are observable in genomes of natural isolates. Theoretical models show that fluctuating environments can enable the persistence of stable TE colonies in asexual and selfing populations if TE provide fitness or evolvability benefits to the host^59,60^. We speculate that the persistence of Tf elements in fission yeast genomes depends on their dynamic contribution to fitness, showing a mutualistic rather than parasitic symbiosis.

While Tf2 insertions can provide a fitness advantage to its host, the negative effects of CoCl_2_ treatment and mating type on the WT strain indicate that they may also constitute a genetic burden that could offset their positive effects. Both CoCl_2_ treatment^24^ and Mc expression^42^ may drive increased Tf2 transcription that could, as detailed above, result in replicative stress^30,50,61^. An experiment in *S cerevisiae* that overdosed its genome with Ty1 copies revealed that they cause a decreased capacity to survive genotoxic insults and inhibitors of DNA replication^62^. A similar mechanism could explain the negative fitness effect of Tf2 transcription increase. The growth rate of natural isolates upon challenge with CoCl_2_^26^ is negatively correlated with TE copy number (Supplementary Figure 5), supporting a model where overall Tf1 and Tf2 dosage decreases fitness when TE expression is induced. Together, our results provide evidence for a model in which Tf1 and Tf2 elements constitute a dynamic source of regulatory variation with positive fitness contribution. In this view, the Tf1 and Tf2 elements exhibit a very low transposition rate, but an increased chance of generating insertions with a positive fitness effect through Sap1 mediated target site selection. Recombination can then both fine-tune their fitness contributions by deletion of insertions in response to changing environmental conditions and maintain persistence through the generation of tandem arrays. More generally, we show that these and other fundamental aspects of TE biology may be directly addressed through the use of transposon free strains.

## Acknowledgements

We are grateful to Henry Levin for valuable discussions and reagents, Jef Boeke for providing Tf2 overexpression constructs, and Megan Behringer and Ashley Farlow for information and discussions regarding MA line experiments. This work was supported by grant R35GM131763 from NIH/NIGMS.

## Author Contributions

SCM, CE and MZ conceived the project, carried out experiments and wrote the manuscript. RP generated the transposition reporters. ER performed the RNAseq experiments on *sap1* mutants. NC measured transposition by Tf2 overexpression in Tf-null and *sap1* mutants. WC and CE performed long-read sequencing and assembly of Tf-null and parental strains.

## Data availability

All High Throughput Sequencing data generated in this work are available in the NCBI/NIH Sequence Read Archive under Bioproject ID PRJNA767600.

## Code availability

The code used to analyze the data in this work will be made available upon request

## Methods

### Analysis of natural isolates sequence data

To reanalyze the sequencing data from Jeffares et al ^26^ we downloaded the raw FASTQ files corresponding to the non-redundant 57 clonal strains identified in this study from the European Nucleotide Archive Accession PRJEB2126. We carried out copy number analysis with the DeviaTE pipeline^28^ on trimmed and quality filtered reads using a database including the divergent region of the gag gene from Tf1 and Tf2 (positions 56-1000 with position 1 being the first nucleotide after the 5’ LTR), and a set of single copy essential genes from regions showing no evidence of amplification or deletion^63^ (myo2, mrpl24, cdc6, urb1, cog2, cca1, spc7, tti1, dna2, lsh1, rpc1, psm1, cut3, msi2, sfc3, cog1, med15, ero12, cdc7, rpb3, alp1, smc5 and sws2). The output files were parsed to retrieve the High Quality estimated copy number. To validate this approach, we plotted the DeviaTE HQ copy number estimates with the copy number detected by BLAST of the same region against long-read derived genome assembles from the strains for which these were available as reported by Tusso et al^39^ (SRA accession PRJNA527756). For insertion site detection and characterization, we first aligned the trimmed and quality filtered reads with BWA v0.7.17 mem against the *S pombe* genome assembly ASM294v1.16, and prepared the resulting bam files with the samtools v1.7 suite programs sort -n, fixmate, sort and index. We carried out two separate MELT v2.2.2^29^ searches on the alignment files using Tf1 and Tf2 consensus sequences and the annotation of Tf1 and Tf2 elements in the type strain obtained from Bowen et al ^18^. We then filtered, combined and consolidated the resulting vcf files to obtain a maximal catalog of potential insertion sites. Using this, we retrieved the paired reads mapping to within 800bp of each insertion site from the individual alignment file showing evidence of a potential insertion, and used them to assemble individual mini-contigs using SPADES v3.12.0 –careful –only-assembler. We then aligned each minicontig generated by SPADES using BLAST 2.2.31 against a database of consensus LTR from Tf1 and Tf2. We parsed the BLAST output to select the top hit for each LTR and select minicontigs with more than 150bp in alignment overlap. Using this information, we then retrieved the sequence from the minicontig flanking the LTR alignment into individual fasta files using seqkit 0.16 grep. We then blasted the sequences flanking the LTR to the *S pombe* genome to identify the insertion sites and generate a maximal list of detected insertions with strain and LTR identification and orientation. This list was manually curated to consolidate and merge insertions detected in the same strain and with the same position and orientation. The few cases of insertions within the same strain and with the same position but different LTR identification or orientation were restricted to strain JB1207, the only diploid in the collection. Insertions with multiple identified LTR or incongruent orientations between the flanking sequence alignments, usually corresponding to nested insertions, were individually realigned to discern the position of the oldest insertion. Inner nested insertions were discarded. We then processed the list of insertions to generate a presence-absence matrix, calculate frequencies in the population, and generate the plots in figure 1.

To calculate the apparent ratio of transposition over deletion in the population we fit a simple transposition/deletion model on the distribution of estimated gag copy number and the total number of insertions present in each strain with at least one copy of Tf1 or Tf2 elements. In this simple model, each new transposition will create an indelible LTR insertion, growing with a rate ΔLTR = μTfX, where TfX is the original number of full length Tf1 or Tf2, and μ is the transposition rate. The full-length elements (as estimated by gag copy number) will change with a rate Δgag=TfX(μ-ν), where ν is the deletion rate. Dividing, Δgag/ΔLTR=(1- ν/μ), and the apparent ratio of transposition to deletion rates μ/ν can be estimated from the slope of the linear regression of gag copy number to total insertions. We note that this model requires complete neutrality to provide a good estimate of the ratio of the real μ and ν, as selection could alter the apparent transposition and deletion rates depending on the average fitness effects of new insertions and their deleted counterparts. We carried out estimation of μ/ν with a model specified in the STAN programming language in the R environment using package rstan.

### Cloning and Constructs

All oligonucleotides used in this study are in Supplementary Table 4. To generate a Tf2 transposition reporter plasmid we cloned a consensus Tf2 into the SmaI digested pBSade6 integration plasmid by Gibson assembly of the 5’LTR, and the coding sequence with the neoR-Artificial Intron (neoRAI) and the 3’ LTR, obtained by amplification with primers oM2382/ oM2386 and oM2385/oM2383 respectively. The LTR and CDS-containing fragments were amplified from plasmids pHL1631 (WT) and pHL1632 (gagfs), a kind gift from Jef Boeke and Henry Levin. The resulting insertion plasmids were pMZ851 (Tf2WT) and pMZ855 (Tf2gagfs). To generate deletion reporters of Tf2-6 with a *ura4* insertion, we first generated a CRISPR plasmid with two gRNA expression cassettes that would cleave both sides of Tf2-6. First, we cloned a single gRNA CRISPR plasmid by Gibson assembly of a fragment amplified from pDB4283^64^ by oligos oM1568/oM1762, with NotI digested pDB4281, resulting in plasmid pMZ677. We then inserted an additional gRNA expression cassette by Gibson Assembly of two PCR products obtained by PCR of pMZ677 with primer pairs om1770/om1771 and om1772/om1773, together with BsrGI digested pMZ677, resulting in pMZ691. Next, we assembled Tf2-6::*ura4*+ HR donors containing flanking LTR, no LTR and a duplication of 350bp present upstream of the 5’ LTR in place of the LTR. First, we assembled pMZ743 by Gibson assembly of a PCR product amplified from pBluescript with oM2235/oM2238, together with PCR products by amplification of strain PEY305 genomic DNA with primer pairs om2237/2239, om1315/2240, om1316/2242, om2241/2236, resulting in pMZ743. We then removed the MCS from the *ura4* insertion in pMZ743 by Gibson assembly of PCR products amplified from pMZ743 with primer pairs om2265/om2266 and om2268/om2267, yielding plasmid pMZ746. We removed the LTR by Gibson assembly of PCR products from pMZ746 with primer pairs oM2281/oM2282 and oM2280/oM2283, yielding pMZ753, which was used as a HR donor to generate Tf2-6::*ura4+* reporters with no LTR. A polymorphism present in the 5’ LTR was removed to make both LTR identical, by reassembling the pMZ746 plasmid with PCR products from primer pairs om2257/om2260 and om2258/om2259, yielding pMZ762. To assemble the Tf2-6::*ura4+* donor with flanking tandem repeats we Gibson assembled PCR products from pMZ753 with primer pairs om2276/om2278 and om2277/om2279, yielding pMZ766. The plasmids for CRISPR-aided Tf2 removal were generated by ligation of phosphorylated and annealed oligonucleotide pairs containing the targeting sequence into CspCI digested pMZ374^65^. The plasmid containing the *ura4* gene flanked by Tf2CDS homology regions (pMZ160) was generated by amplification of flanking homology regions of Tf2 from strain 972 genomic DNA with primer pairs Tfam_USF/Tfam_USR-ura4 and Tfam_DSF-ura4/Tfam_DSR. The fragments were then used in a megaprimer PCR reaction with plasmid pUR19, containing the *ura4* gene, and primers Tfam_USF/Tfam_DSR to generate a fragment that was then digested with XhoI and cloned into XhoI digested pUC19. For deletion of Tf2- fragment1 (which lacks the homology in the right arm of the fragment from pMZ160) we amplified the *ura4* gene with Tf2 homology arms from plasmid pMZ762 with primers oM1781/oM1225.

To generate the *sap1-c* mutation with CRISPR, we cloned the targeting sequence into pMZ374 as above using oligos oM1172/oM1173, generating plasmid pMZ544.

### Strains growth and genetic manipulation

All strains used in this study are detailed in Supplementary Table 5. Media recipes were as described in the Nurse Laboratory Handbook. All transformations were carried out using the Lithium Acetate/PEG heat shock method.

Deletion reporters were generated by transformation of strains PB1 (h90) and CHP429 (h-), or ZB2952 (h+Tf-null) or ZB2950 (h-Tf-null) with double-gRNA CRISPR plasmid pMZ691 and the HR donor from plasmids pMZ766, pMZ753 or pMZ762 isolated by KpnI/SalI digestion followed by agarose gel electrophoresis and band extraction. Transformed cells were plated on YEA-Blasticidin (30mg/L) plates, followed by replica plating onto EMMG+dropout supplements without uracil with 30mg/L Blasticidin to select transformants that incorporated the *ura4* marker.

Candidate strains were validated by PCR-genotyping, sequencing over the Tf2-6 LTRs and southern blotting to rule out tandem repeat insertions, and back-crossed to ensure that a single *ura4* insertion had occurred.

To generate transposition reporters the plasmids pMZ854 and pMZ855 were digested with ZraI, transformed into strains ZB3147 and ZB3142, and selected in media without adenine, to obtain the transposition reporters in WT and Tf-null backgrounds respectively.

To generate the *sap1-c* mutation by CRISPR we transformed strains PB1 and ZB1925 with plasmid pMZ544 together with a fragment obtained by annealing of oligonucleotides oM1170/oM1171 as a homologous recombination sequence donor, and plated on media without uracil. Survivors were screened for the presence of the *sap1-c* mutation by colony PCR with primers oM6/oM7 followed by digestion with BclI.

### Transposition and deletion assays

To measure transposition, prototrophic strains with the indicated Tf2WT and Tf2gagfs transposition reporters were grown in octuplicate cultures in EMM to saturation, and then plated at a density of 5e9 cells per plate in YE-G418 plates and 5e2 per plate in YE. The frequency of G418 positive cells was fit to a maximum likelihood fluctuation model to calculate the transposition rate per generation. Colonies from the YE-G418 plates were characterized by PCR with primers oM146 and one of primers oM1366-1377 (Tf2-1 to Tf2-13), oM2049 (Tf2-8), oM2210 (Tf2- frag1) or oM2224 (Tf2-14) to identify insertions by GC into entopic Tf2 elements. Auto-GC (loss of artificial intron in the neoRAI reporter by GC with cDNA) was ruled out by way of a PCR amplifying the neoR gene from the inserted plasmid (primer pair oM179/oM1761) followed by digestion with SpeI, which has a restriction site in the artificial intron that would be removed through GC with cDNA. All G418 resistant colonies showed the presence of the SpeI site, indicating that auto-GC had not occurred.

To measure deletion rates, we grew the indicated Tf2-6::*ura4*+ strains in YNB media and then plated cells at a density of ∼1e7 in YNB-5FOA and ∼1e3 in YNB+Dropout complete. We then fit the frequency of *ura4* loss to a maximum likelihood fluctuation model to calculate *ura4*-loss rates. Colonies from the YNB-5FOA plates were characterized by PCR with primers oM1365/oM1371/oM2038 and oM

To measure transposition by overexpression of Tf2, we transformed strain ZB1925 (CRISPR-derived Tf-null) and PB1 (WT) with inducible expression plasmids pHL1631 (Tf2), pHL1632 (Tf2gagfs) and pHL1633 (Tf2INTfs), which drive overexpression of Tf2 from the nmt1 promoter inducible in the absence of thiamine. Four independent colonies of transformed cells were patched onto EMM plates lacking thiamine and grown for 4 days at 32**°**C. After, cells were patched onto 5-FOA to remove the Tf1-*neo* expression vector, and dilutions were plated onto 5-FOA or YES+G418+FOA+2g/L drop-out minus uracil mix plates to measure transposition frequencies. The proportion of G418 resistant colonies to 5-FOA resistant colonies represents the transposition frequency.

### Analysis of MA strains

The sequences from the Farlow et al^38^ and Behringer et al^37^ MA studies were downloaded from the SRA (Accessions PRJNA295384 and PRJNA301358 respectively). For new insertion detection we used the same MELT/minicontig analysis pipeline described above. For copy number estimation we used the same DeviaTE pipeline described above. All copies in the Tf2-7/8 array exhibit the exclusive T2250G polymorphism (with position 1 being the first nucleotide after the 5’ LTR). We parsed the DeviaTE output to obtain the frequency of the T2250G polymorphism and estimated the size of the Tf2-7/8 array and the copy number of originally monomeric insertions. To estimate the rates of ICR and USCE from the distribution copy number of monomeric (Not Tf2-7/8) insertions, we considered a model in which ICR events cause a loss of -1 copy with a rate γ per generation, and USCE events cause either a gain of +1 copy or a loss of -1 copy with equal probability of each, with a rate of θ per generation. Assuming independence between different Tf2 elements this can be modeled as a biased 1-dimensional random walk with a boundary condition at 0. By simulation we observed that the boundary condition can be safely ignored as long as γ and θ are smaller than 1/(number of Tf2xNumber of generations), in this case, ∼5e-5. Without the boundary condition the rate of ICR contributes to both the mean and the variance of the distribution because of its net negative effect, and the rate of USCE contributes only to the variance because of its net neutral effect. The expected value and the variance of the total copy number of monomeric Tf2 at generation g can be expressed as:

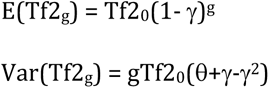

Where Tf2_0_ is the original number of monomeric Tf2. From these two expressions and the observed mean and variance of the distribution we can calculate the rates of ICR and USCE affecting monomeric Tf2 insertions. We carried out estimation of γ and θ and their standard error using R with the bootstrapping package boot.

### Generation of Tf-null strains

The relatively low number of Tf2 insertions in the laboratory type strain makes deletion of all TE in the fission yeast genome uniquely feasible in contrast to other eukaryotic model organisms. In order to obtain rapid TE removal we leveraged the high frequency (∼1e-5) of Tf2 CDS deletion by ICR and USCE. By applying negative selective pressure to the presence of the CDS it should be possible to progressively remove all Tf2 CDS, leaving solo LTR in their stead.

We first attempted to select for Tf2 deletion by directing CRISPR-mediated double strand breaks using a series of CDS-targeted single guide RNAs^65^. We reasoned that cleavage would destabilize the insertion, engaging Homologous Recombination, and provide a negative selection for the presence of the CDS that would enable us to isolate deletion events. We designed and cloned single-plasmid gRNA/Cas9 expression vectors directed at multiple target sites present in the CDS (cloning detailed above) and equipped with a negative/positive selection marker (*ura4*) that enabled rapid cycles of CRISPR (Supplemental Figure 5a and b).

For CRISPR aided Tf2 removal, we transformed strain PB1 with the plasmids detailed in Supplementary Figure 5c and plated in media lacking uracil. Survivors were genotyped for the presence or absence of each entopic Tf2 by colony PCR with combinations of oM1365 with one of oM1366-1377. Survivors with multiple Tf2 deletions were selected, streaked on 5-FOA media to remove the CRISPR plasmid, and retransformed with a different plasmid. The process was repeated until all Tf2 elements were deleted.

This strategy yielded two independent strains showing complete loss of Tf2 CDS after four rounds of CRISPR plasmid transformation and removal (Supplemental Figure 5c). However, these strains exhibited very poor spore viability upon back- cross with the original strain, but not in a selfing cross, suggesting that they had acquired chromosomal rearrangements^66^. Indeed, long-read high throughput sequencing revealed multiple rearrangements, including pericentric and paracentric inversions, balanced translocations and deletions (Supplemental Figure 5d and 5e). The breakpoints of these rearrangements were the Tf2 insertions targeted by CRISPR. Characterization of the partially deleted intermediate strains revealed that the rearrangements could occur without deletion of the CDS, in a transformation round previous to the one that caused deletion by inter-LTR recombination. Thus, while CRISPR/Cas9 cleavage does provide a rapid method to select for inter-LTR recombination, it also leads to non-allelic recombination between the interspersed Tf2 insertions.

We then undertook a classic recombineering approach to remove Tf2 CDS (Supplemental Figure 5f). By transformation of a linear DNA fragment consisting of the *ura4* gene flanked by Tf2 CDS homology arms (Supplemental Figure 5g, cloning detailed above) we were able to randomly tag individual Tf2 insertions through selection in media without uracil. Subsequent selection in 5FOA media yields strains where the *ura4* gene has been lost either by GC, reverting the insertion to its native state, or by inter-LTR recombination, which removes the CDS leaving a solo LTR. We then selected strains with deleted individual Tf2 CDS and combined the deletions by crossing. This process can be repeated until all Tf2 CDS are removed (Supplemental Figure 5h).

For *ura4*-aided Tf2 removal, strains CHP428 and CHP429 were transformed with XhoI digested pMZ160 and plated in media without uracil. Colonies were then genotyped for the position of the *ura4* gene by colony PCRs with primer oM1968 and one of the upstream primers specific for each entopic Tf2 copy (primers oM2033 to oM2044). Colonies with an unambiguously localized *ura4* insertion in an identified Tf2 entopic copy were grown to saturation in liquid YEA and spread onto 5-FOA media plates to select *ura4* loss events. The surviving colonies were genotyped for deletion of the Tf2 copy that received the *ura4* insertion by colony PCR with primer oM1365 and an upstream/downstream primer pair corresponding to the Tf2 copy of interest (Downstream primers: oM1366-oM1377; Upstream primers: oM2033-oM2044). Candidates with deletions were genotyped for mating type with primers oM9/oM10/oM11 and for ade6-M210/M216 allele with primers oM12/oM13 followed by XhoI digestion. Once classified in h- and h+ groups, the deletions were combined by crossing between mating-compatible strains and genotyped for segregation of the deleted allele as above. Segregants with the desired combinations were used for a subsequent cycle of Tf2 removal. In the process of removing all described Tf2 entopic copies we detected a new Tf2 insertion present in the CHP428/CHP429 background, located in coordinates I:4940032, which we then deleted in the same manner as the rest. The complete genealogy of the *ura4*- aided Tf2 removal is detailed in Supplementary Figure 5h.

We performed long and short read whole genome sequencing to fully characterize the genome of the original parental strains and the Tf-null derivatives. The assembled genomes confirmed the complete removal of all Tf2 coding sequences with no rearrangements or deletions (Supplemental Figure 5I). The strains obtained from the Tf2-CDS removal process were isogenic with the parental strains and had only acquired four single nucleotide polymorphisms, of which only one was missense, located in non-conserved region of a protein coding gene (Supplementary Table 2).

The Tf-null strains ZB2950/ZB2952 and the parental strains CHP428/CHP429 were cured of the present auxotrophies (*ade6-M210* or *ade6-M216*, *leu1-32*, *his7-366* and *ura4-D18*) to generate prototrophic strains by serial transformation with PCR fragments containing the WT alleles (primer pairs: oM12/oM13 – ade6-M210; oM2300/oM2301 – leu1-32; oM2302/oM2303 – his7-366, oM2304/oM2305 – *ura4*-D18) followed by selection in EMM media without the corresponding supplement and confirmation by Sanger sequencing, resulting in strains ZB3152,ZB3153 (from CHP428 and CHP429 respectively) and ZB3154,ZB3155 (from ZB2952 and ZB2950 respectively)

### Competitive growth assays

For competitive growth assays, pairs of otherwise isogenic h+ or h- Tf-null and WT prototrophic strains (ZB3152/ZB3154 and ZB3153/ZB3155) were grown separately in liquid EMMN to exponential phase, harvested and washed twice in EMMN, counted and combined in equal numbers to an OD of 1, and then distributed to triplicate 5ml cultures of EMM or EMM +250μM CoCl_2_ at an OD of 0.025. 1 OD of the initial mix was harvested by centrifugation and frozen as Generation 0. The cultures were serially transferred to 5ml of fresh media twice a day to OD 0.1 in the morning and OD 0.025 at night, to prevent them from growing past OD 3. We measured OD at every passage to keep a record of the generations passed. Every 2 days approximately 1.5 OD of each culture were harvested by centrifugation and frozen. After 8 timepoints had been harvested (14 days, mean=136.4 generations and SD=0.55 for EMMN, mean=124.8 generations and SD=0.52 for EMMN+250μM CoCl_2_) we stopped the experiment. We carried out the competition assay on saturation conditions similarly: pairs of h+ and h- Tf-null and WT prototrophic strains were grown together as before, but cultures were passaged every 48 hours to allow them to reach saturation: the cultures reached OD 10 within 24 hours and remained at that OD until passage into fresh media to OD 0.025. We carried out the experiment for 14 days for a total of 60 generations.

To quantify the proportion of genotypes in the harvested competition cultures, we purified the genomic DNA from each timepoint with the Monarch gDNA extraction Kit (New England Biolabs). We then amplified the region of one of the mutations acquired by the Tf-null strain (I:1453108 C->A) with primers amenable to Amplicon-seq with in-line barcodes to identify the samples in pooled sequencing runs (oM2327-oM2339), purified the PCR products, measured their concentrations and pooled them in groups of 12 for Amplicon-Seq (Genewiz, Piscataway, NJ). We processed the resulting FASTQ files removing sequences shorter than 100nt, followed by splitting with fastx_barcode_splitter with options --bol –exact, and then trimming with fastx_trimmer -f 59 -l 63 -Q33 and then collapsing with fastx_collapser -Q33. The output files were parsed to separate the counts for the WT and Tf-null genotypes.

We analyzed the data fitting a Bayesian model specified in the STAN programming language in the R environment using package rstan. In this model^43^, the proportion of Tf-null genotypes in the competition:

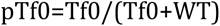

progresses at each timepoint t, separated by Δg generations as:

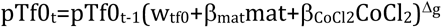

Where w_tf0_ is the relative fitness of the Tf-null/h+ strain in EMMN media with respect to the WT/h+ strain, mat and CoCl_2_ are 0|1 variables representing the h- genotype and CoCl_2_ 250μM treatment, β_CoCl2_ and β_mat_ are coefficients representing the effect of CoCl_2_ treatment and h- genotype.

### Gene expression analysis

To study gene expression differences between the WT and Tf-null strains we grew triplicate 50ml EMMN cultures of ZB3153 and ZB3155 in the presence and absence of CoCl_2_ 250μM from an OD 0.025 to OD 1. We then harvested the cultures, washed them in ice cold water and snap froze the pellets in a dry ice/ethanol bath. We purified total RNA with hot acid phenol extraction in TE +1% SDS, followed by two phenol:Isoamyl alcohol extractions and one Chloroform:Isoamyl alcohol extraction and precipitation with 250mM NaCl and 3 volumes of Ethanol. We air-dried the pellets and resuspended them in DEPC treated water. We treated 100ug of total RNA with the DNAfree DNA removal system (ThermoFisher) and submitted for strand- specific total RNAseq to Genewiz (Piscataway, NJ). For analysis of ncRNA expression in WT (972) and *sap1-c* (ZB973) strains we seeded from EMMN cultures grown overnight into YEA at OD 0.025 and harvested the cultures at OD 0.5 (early exponential growth) and OD 3 (early saturation). RNA was extracted and processed as above.

The FASTQ reads were filtered and trimmed with Trimmomatic 0.32, mapped to the gtf annotation file of the EMSEMBL genomes ASM294v2 assembly with TopHat 2.1.1 with options -r 200 --library-type fr-firststrand, followed by the HTSeq v1.12.4 framework HTSeq.scripts.count -s reverse to assign counts to features. The count data of the CoCl_2_/Tf-null experiment was analyzed with R package DEseq2. The expression of intergenic ncRNA (annotation retrieved from pombase.org/query with product type: feature_type_ncRNA_gene and then filtered to retain only intergenic ncRNA) in the *sap1-c*/growth stage experiment was analyzed with CuffLinks/CuffDiff and R packages lmer4 and emmeans. ncRNA were classified as Sap1 associated if they were within 500bp of a significant Sap1 summit as assessed by MACS analysis. Functional enrichment analysis was carried out on protein coding genes from the EMSEMBL genomes ASM294v2 assembly annotation as assigned by closest proximity to Sap1-associated ncRNA, using the AnGeLi^67^ Web Interface (http://bahlerweb.cs.ucl.ac.uk/cgi-bin/GLA/GLA_input).

---

**Supplementary Figure 1.**
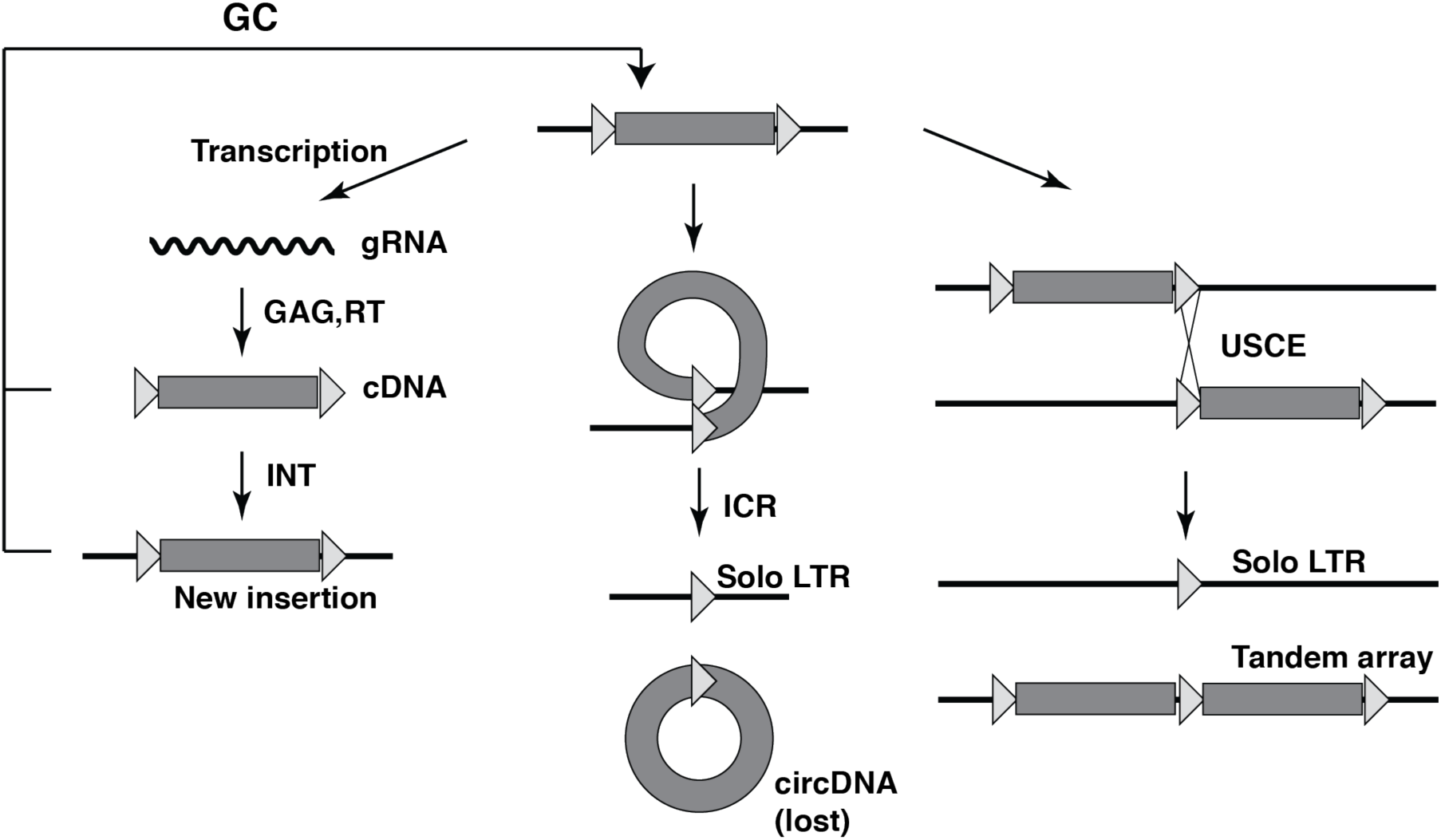
Recombination and Transposition pathways for LTR Retrotransposons. RT: Reverse Transcriptase. INT: Integrase. gRNA: genomic RNA. cDNA: complementary DNA. GC: Gene Conversion. ICR: Intra-Chromatic Recombination. USCE: Unequal Sister Chromatic Exchange.

**Supplementary Figure 2.**
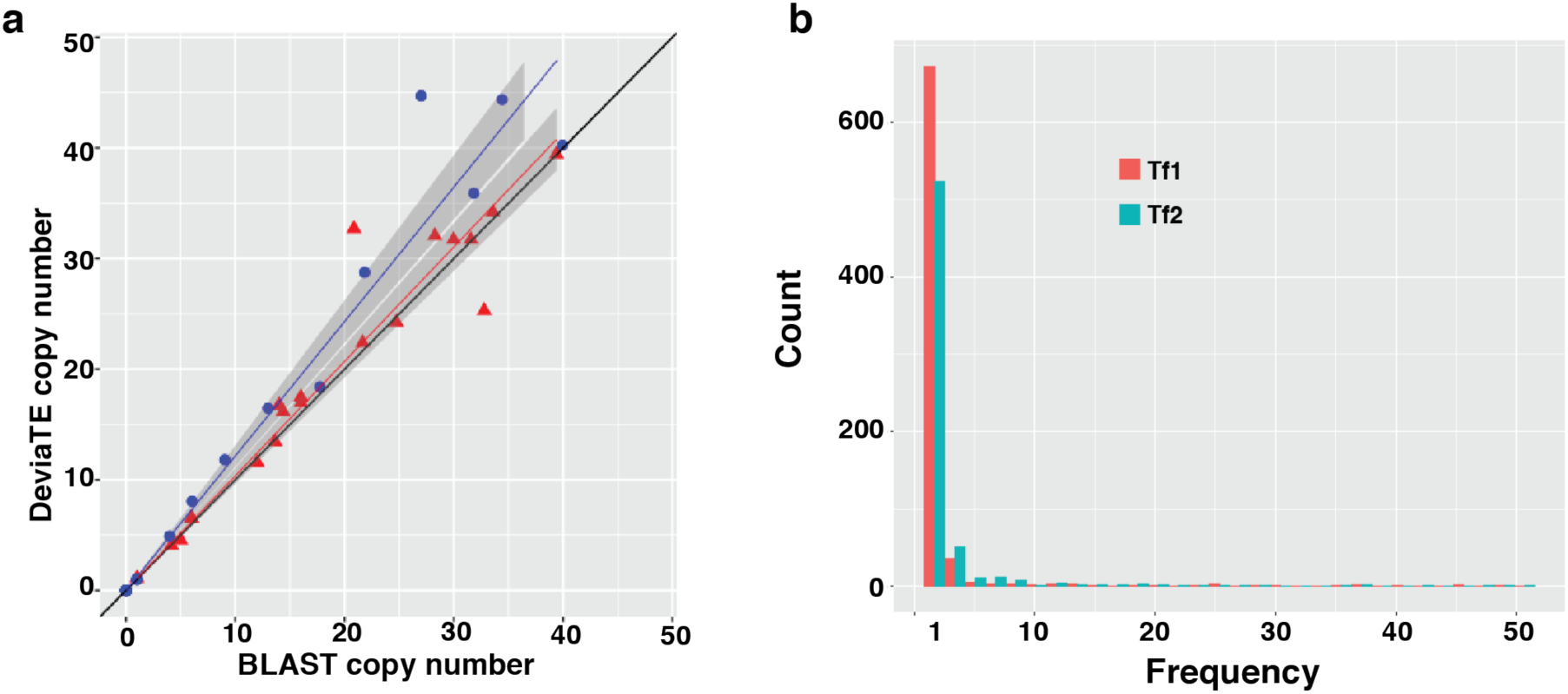
a. Validation of DeviaTE copy number estimation from short read sequencing and BLAST copy number estimation from Long Read assemblies. b. Allele frequency spectrum of polymorphic Tf1 and Tf2 insertions in the 57 natural isolates in Jeffares et al 2015.

**Supplementary Figure 3.**
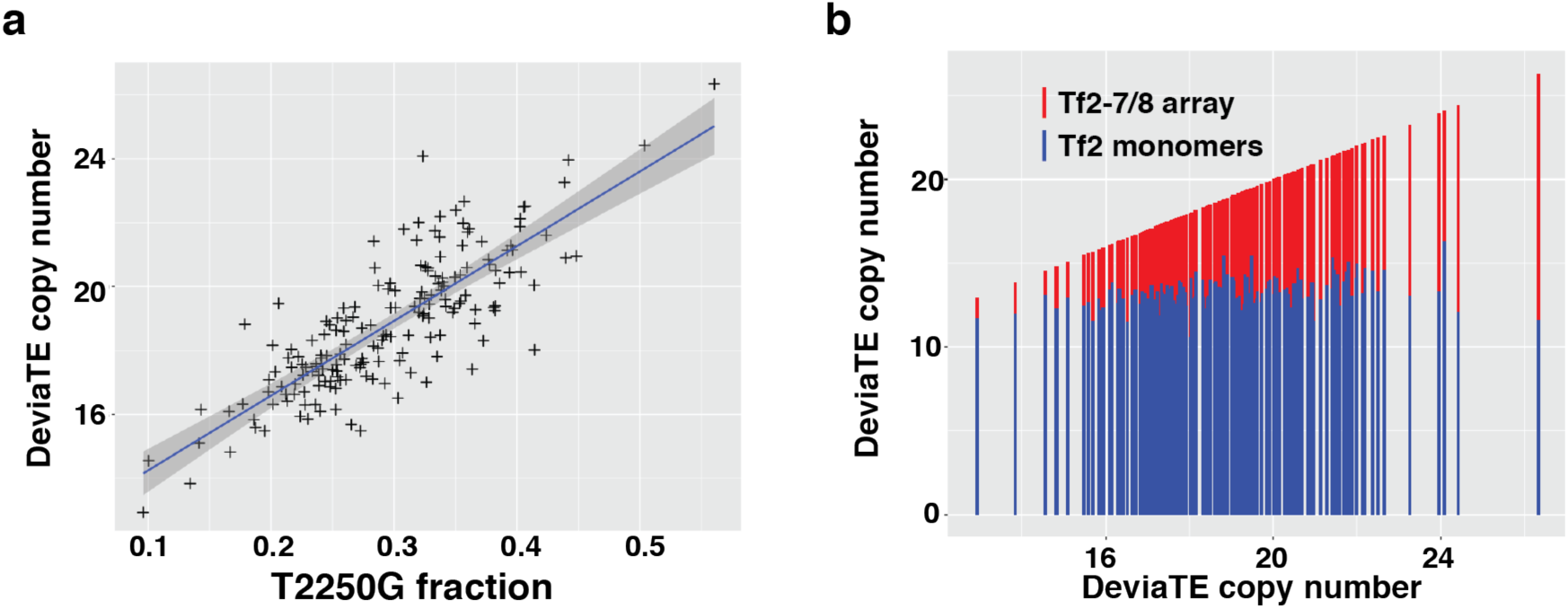
Copy number analysis in MA lines. a: Scatterplot of proportion of reads with the T2250G polymorphism (Tf2-7/8) present over total coverage with Total estimated Tf2 copy number. Line represents best linear regression fit. b: Bar chart of total Tf2 copy number divided by estimated Tf2-7/8 copy number (red) and monomeric Tf2 copy number (blue), arranged by increasing total Tf2 copy number.

**Supplementary Figure 4:**
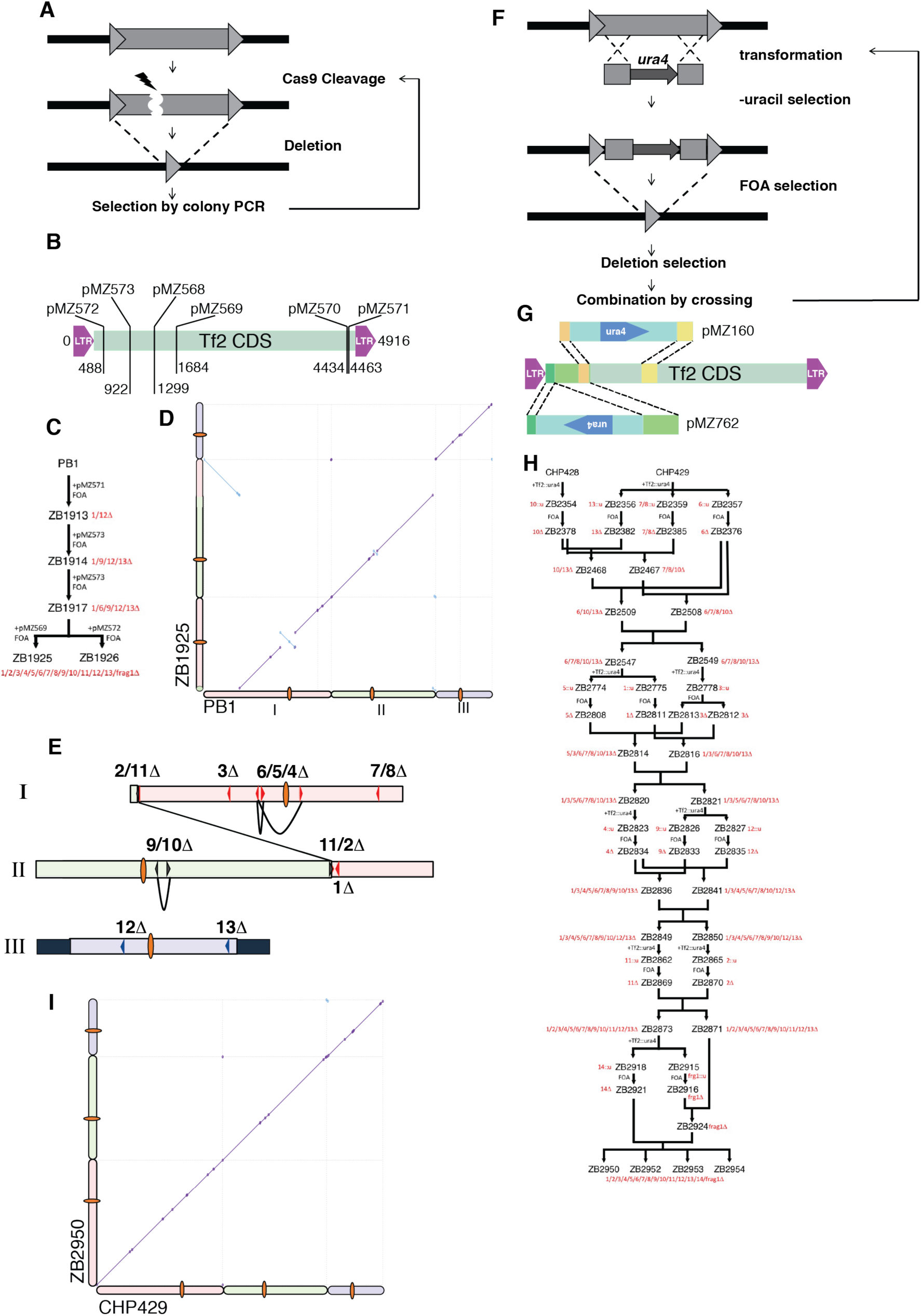
Generation of Tf-null strains by Tf2 transposon removal. a. CRISPR-aided ICR removal method. b. Map of gRNA targets over Tf2 CDS with CRISPR plasmid names. c. Genealogy of CRISPR-derived Tf-null strains. d. Genome-wide dot-plot alignment for a CRISPR-generated Tf-null strain (ZB1925) and the parental strain (PB1). e. Diagram of rearrangements detected in panel d. f. Classical recombineering ICR removal method. g. Schematic of Tf2::*ura4* transforming fragments used in the recombineering Tf2 deletion method. h. Genealogy of Recombineering-derived Tf-null strains. i. Genome-wide dot-plot alignment for a recombineering-generated Tf-null strain (ZB2950) and the parental strain (CHP429).

**Supplementary Figure 5.**
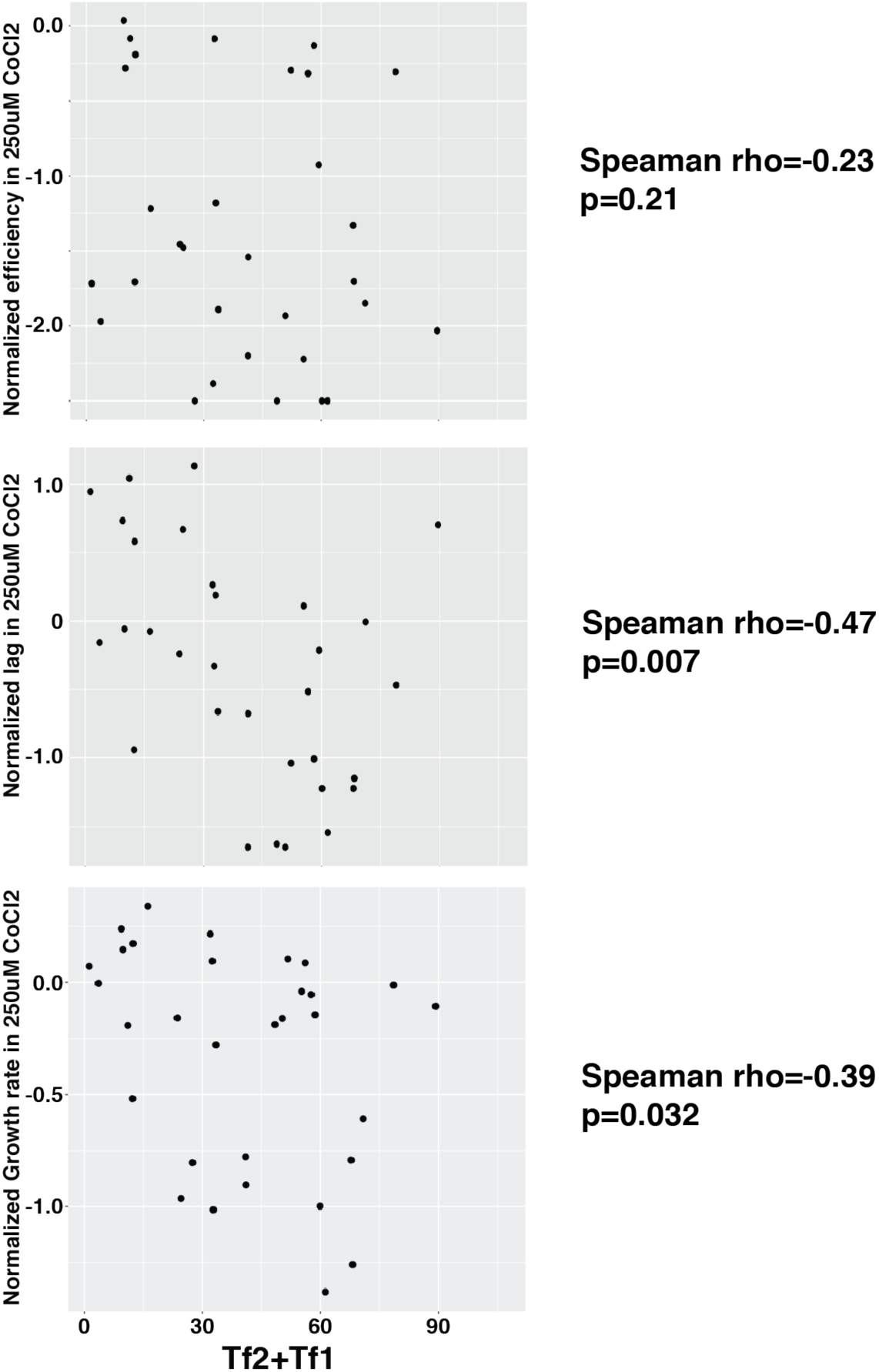
Scatterplot of growth parameters in the presence of CoCl_2_ of natural isolates described in Brown et al 2011.

